# From people to *Panthera*: Natural SARS-CoV-2 infection in tigers and lions at the Bronx Zoo

**DOI:** 10.1101/2020.07.22.213959

**Authors:** Denise McAloose, Melissa Laverack, Leyi Wang, Mary Lea Killian, Leonardo C. Caserta, Fangfeng Yuan, Patrick K. Mitchell, Krista Queen, Matthew R. Mauldin, Brittany D. Cronk, Susan L. Bartlett, John M. Sykes, Stephanie Zec, Tracy Stokol, Karen Ingerman, Martha A. Delaney, Richard Fredrickson, Marina Ivančić, Melinda Jenkins-Moore, Katie Mozingo, Kerrie Franzen, Nichole Hines Bergeson, Laura Goodman, Haibin Wang, Ying Fang, Colleen Olmstead, Colleen McCann, Patrick Thomas, Erin Goodrich, François Elvinger, David C. Smith, Suxiang Tong, Sally Slavinski, Paul P. Calle, Karen Terio, Mia Kim Torchetti, Diego G. Diel

## Abstract

We describe the first cases of natural SARS-CoV-2 infection detected in animals in the United States. In March 2020, four tigers and three lions at the Bronx Zoo developed mild respiratory signs. SARS-CoV-2 RNA was detected by rRT-PCR in respiratory secretions and/or feces from all seven affected animals; viral RNA and/or antibodies were detected in their keepers. SARS-CoV-2 was isolated from respiratory secretions or feces from three affected animals; *in situ* hybridization co-localized viral RNA with cellular damage. Whole genome sequence and haplotype network analyses showed tigers and lions were infected with two different SARS-CoV-2 strains, suggesting independent viral introductions. The source of SARS-CoV-2 infection in the lions is unknown. Epidemiological data and genetic similarities between keeper and tiger viruses indicate human to animal transmission.

---

COVID-19, a severe respiratory disease caused by a novel coronavirus, SARS-CoV-2 (severe acute respiratory syndrome coronavirus-2) (*1*), was first reported in Wuhan, Hubei province, China at the end of December 2019 (*2*). Within weeks the virus caused a global pandemic, and by July, 2020, over 10 million people were infected and more than 500,000 had died (https://www.who.int/emergencies/diseases/novel-coronavirus-2019; accessed 7/1/2020). An early cluster of human cases had an epidemiological link to the Huanan Seafood Wholesale market in Wuhan where a variety of live wild animals were sold (*3*). Genome sequence analysis revealed SARS-CoV-2 to be most closely related to a bat coronavirus (RaTG13-2013), and bats are considered the likely source of the ancestral virus from which the currently circulating SARS-CoV-2 virus was derived (*3*, *4*). Subsequent genetic adaptation in an intermediate animal host(s) or after human transmission has been proposed (*4*, *5*).

Given the suspected zoonotic origin of SARS-CoV-2, identifying susceptible animal species, reservoirs, and transmission routes are topics of global scientific and public interest. Natural SARS-CoV-2 infections in animals have been reported in dogs, cats and farmed mink in Hong Kong, Europe, China and the United States (*6*–*8*). Infection in most of these cases has been linked to households or settings in which human owners or caretakers have tested positive for SARS-CoV-2. Experimental inoculation studies have shown that SARS-CoV-2 infects and replicates with high efficiency in domestic cats, ferrets and fruit bats and poorly in dogs, pigs, and chickens; ducks do not seem to support productive SARS-CoV-2 infection (*9*, *10*). Importantly, virus shedding and horizontal transmission have been shown in cats and ferrets (*9,10*)following experimental inoculation.

In this study, we report natural infection of tigers (*Panthera tigris*), lions (*P. leo*) and keepers who provided care for the animals with SARS-CoV-2 at the Wildlife Conservation Society’s (WCS) Bronx Zoo, New York NY. We provide a detailed characterization of viruses obtained from infected animals and keepers who had close contact with the SARS-CoV-2 positive animals. The animals were infected in March 2020, when, due to widespread community transmission (*11*), NY had become a global SARS-CoV-2 epicenter.

On March 27, 2020, a 4-year-old, female Malayan tiger (*P. t. jacksoni*) (Tiger 1) developed an intermittent cough and audible wheezing despite remaining eupneic. By April 2, an additional Malayan (Tiger 2) and two Amur tigers (*P. t. altaica*) (Tigers 3, 4) housed in the same building as Tiger 1 but in different enclosures, and three African lions (*P. l. krugeri*) (Lions 1, 2, 3) housed in a separate building developed similar respiratory signs. All animals otherwise exhibited normal behavior and activity. Clinical respiratory signs resolved within one week in all but Tiger 1, whose clinical signs lasted 16 days. An additional Amur tiger (Tiger 5) in the same building as Tigers 1-4 did not develop clinical respiratory disease.

Clinical evaluation and a broad diagnostic investigation were performed in Tiger 1. Thoracic radiography revealed small, multifocal regions of peribronchial consolidation. Cytologic examination of tracheal wash fluid identified necrotic epithelial and inflammatory cells consistent with tracheitis (**Fig. 1A** and **B**). *In situ* hybridization (ISH) co-localized SARS-CoV-2 RNA within necrotic epithelial and inflammatory cells in tracheal wash fluid (**Fig. 1C, D and S1**). Targeted PCR testing and metagenomic analysis on respiratory samples (nasal, orpharyngeal, tracheal) were negative for common feline pathogens (**Table S1**, BioProject accession number <u>PRJNA627354</u>). All samples were positive for SARS-CoV-2 by real-time reverse transcription PCR (rRT-PCR) using CDC 2019-nCoV primers and probes targeting the nucleocapsid gene (N1, N2 and N3) and an in-house rRT-PCR assay targeting the envelope (E) gene (**Tables S2** and **S3**). Results were confirmed with MinION and Sanger-based sequencing of the full SARS-CoV-2 spike (S) gene, an internal region of the N gene, and the RNA-dependent RNA polymerase (RdRp) confirmed the virus in the respiratory samples (**Table S2** and **S5**).

**Fig. 1.**
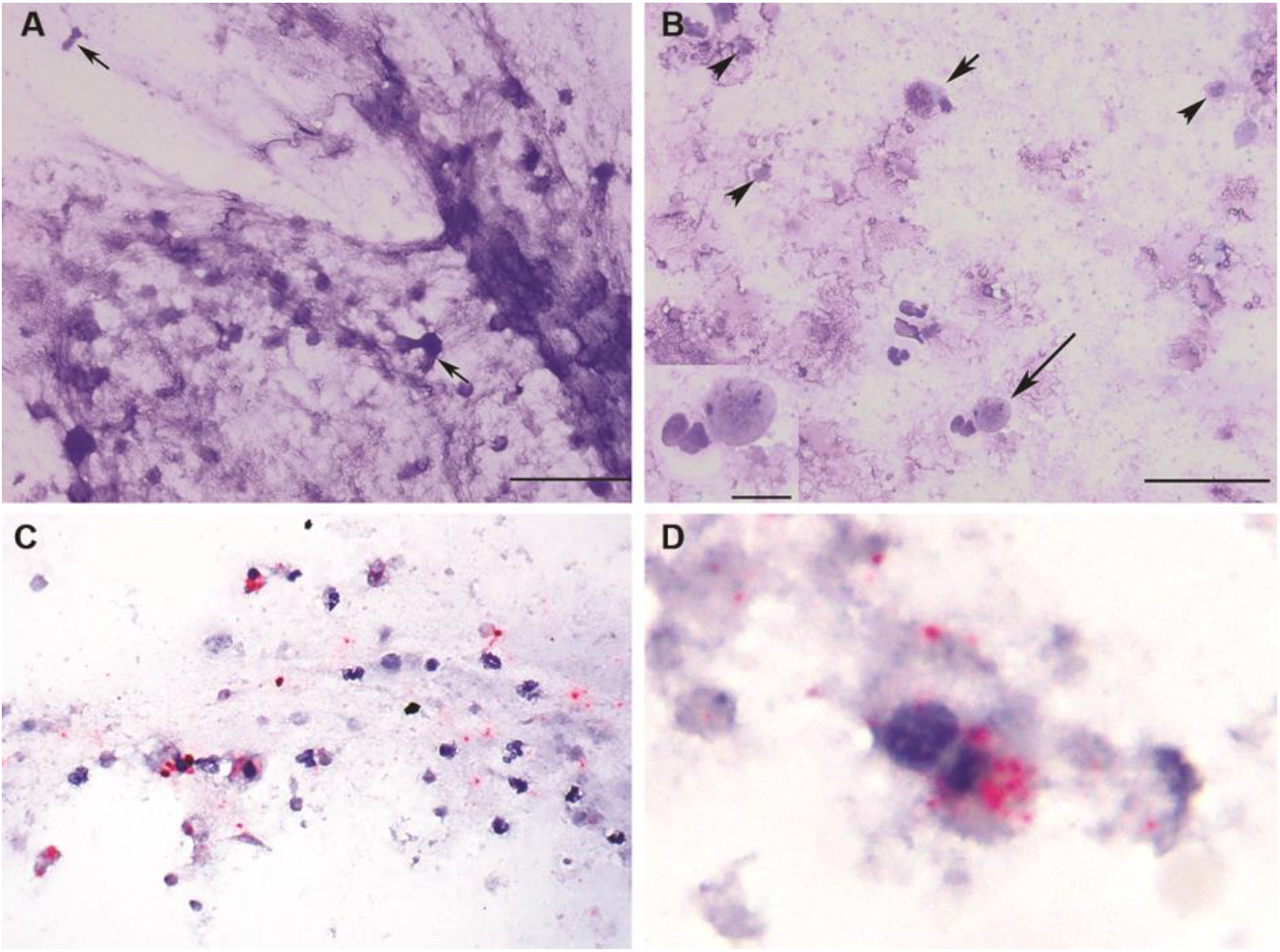
Tracheal wash cytology (A, B) and *in situ* hybridization (ISH) (C, D) in a SARS-CoV-2 infected tiger (Tiger 1). (A) Flocculent material consists of stringy mucus with enmeshed necrotic cells (arrows). Scale = 50 um. (B) Intact columnar (short arrow) and round to cuboidal necrotic epithelial cells with pyknotic nuclei are admixed with neutrophils, macrophages,lymphocytes and necrotic debris (arrowheads). Scale = 50 um. Inset: Epithelial cell with a pyknotic nucleus and a neutrophil. Scale = 10 um. Modified Wright’s-stain (MR). (C, D) probe targeting SARS-CoV2 Incubation with SARS-CoV-2 specific probe shows multifocal red puncta (viral RNA) throughout the mucinous material (blue) and within the cytoplasm of some intact and degenerate epithelial and inflammatory cells. (D) Clumped, degenerate epithelial cells with cytoplasmic red puncta. Surrounding cellular debris contains similar red puncta. Red chromogenic assay with hematoxylin counterstain.

Fecal samples collected opportunistically from each animal (symptomatic Tigers 1-4, asymptomatic Tiger 5, Lions 1-3) were test positive for SARS-CoV-2 by rRT-PCR (**Tables S4**). Results were confirmed by amplicon sequencing (**Table S5**).

Virus isolation was performed in respiratory and/or fecal samples from the animals. Cytopathic effect (CPE) was observed in Vero cells inoculated with tracheal wash fluid from Tiger 1 (**Fig. 2A** and **B**) and fecal samples from Tiger 3 and Lion 2 (**Table S6**). Results were confirmed by rRT-PCR (CDC N1 assay) and/or ISH and immunofluorescence assays (**Fig. 2C** and **D**). A neutralizing antibody titer of 64 detected in Tiger 1 (**Table S7**) confirmed infection and development of an immunologic response against SARS-CoV-2.

**Fig. 2.**
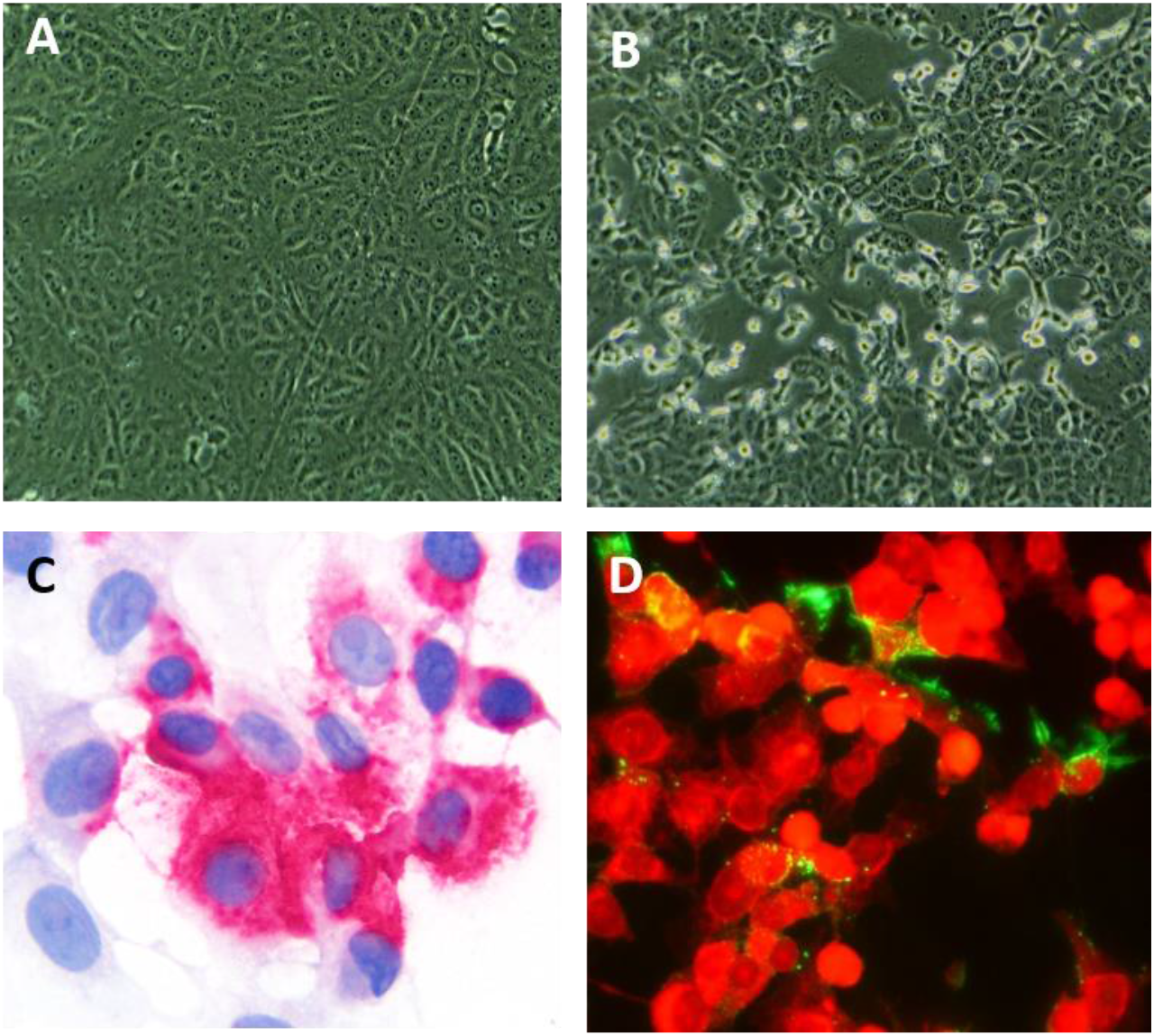
Isolation of SARS-CoV-2 from respiratory specimens from a tiger (Tiger 1). (A) Mock infected control Vero cells and (B) Vero cells inoculated with tracheal wash fluid from the tiger showing typical CPE at 48 h post-inoculation. (C) ISH using a SARS-CoV-2 S-specific probe shows cytoplasmic red puncta in Vero cells at 48 h post-inoculation. Red chromogenic assay with hematoxylin counterstain. (D) Immunofluorescence assay using a SARS-CoV N-specific monoclonal antibody showing replication of SARS-CoV-2 in inoculated Vero cells (green). Evans blue counterstain (red).

An epidemiologic investigation of zoo staff identified ten zoo keepers and two managers who provided care for and had close but not direct contact with the tigers or lions between March 16, 2020 (the date the zoo was closed to the public due to the pandemic) and March 27 to April 1, 2020 (timeline of disease onset in the animals). Four staff (2 tiger and 2 lion keepers) reported mild respiratory symptoms (including fever, cough, chills, myalgia and fatigue) between March 20 and 28, 2020. Oropharyngeal samples and blood were collected from these staff members on April 6, 2020 and tested by rRT-PCR and a microsphere immunoassay (MIA) to detect IgG antibodies. The remaining staff reported no symptoms and were not tested. All tested keepers had evidence of current or prior SARS-CoV-2 infection (one rRT-PCR positive tiger keeper [Keeper 1], one rRT-PCR and serologically positive tiger keeper [Keeper 2], two serologically positive lion keepers [Keepers 3 and 4]). All reported staying at home while sick. Whole genome sequencing of rRT-PCR positive samples from Keepers 1 and 2 was performed to characterize the human samples and further compare the human and animal viral genome sequences.

Nine complete SARS-CoV-2 genome sequences (four from tigers, three from lions, two from keepers) and eight full-length S gene sequences (seven symptomatic and one asymptomatic animals) were generated directly from respiratory and/or fecal samples. Compared to the Wuhan-Hu-1 (GenBank accession number NC_045512), all sequences obtained from the tigers and keepers contained six single-nucleotide polymorphisms (SNPs) with nine additional ambiguous sites (**Fig. S2, Table S8**). A total of 20 sites differed between the three lion sequences and Wuhan-Hu-1 (**Fig. S2, Table S8**).

Viral sequences in the tigers, lions and keepers clustered into common SARS-CoV-2 clades (**Fig. 3A**). Those from tigers and tiger keepers clustered with clade G (defined by the D614G substitution in the spike protein); the lion sequences clustered with clade V (defined by the G251V substitution in ORF3a) (**Fig. 3A**). Median-joining haplotype network analysis of the viral sequences in the animals and keepers corroborated results of the phylogenetic analyses (**Fig. 3B**). Genomic and epidemiological data support a close evolutionary relationship between the viral strains recovered from tigers and tiger keepers. Notably, the genetic differences and the distant phylogenetic relationship between sequences recovered from the tigers/tiger keepers and lions, and the relationship of these strains in the context of global sequences (**Fig. 3B**) indicate that tigers and lions were infected by two different viral strains. Furthermore, this data suggest that at least two independent SARS-CoV-2 introductions occurred, one in tigers and another in lions.

**Fig. 3.**
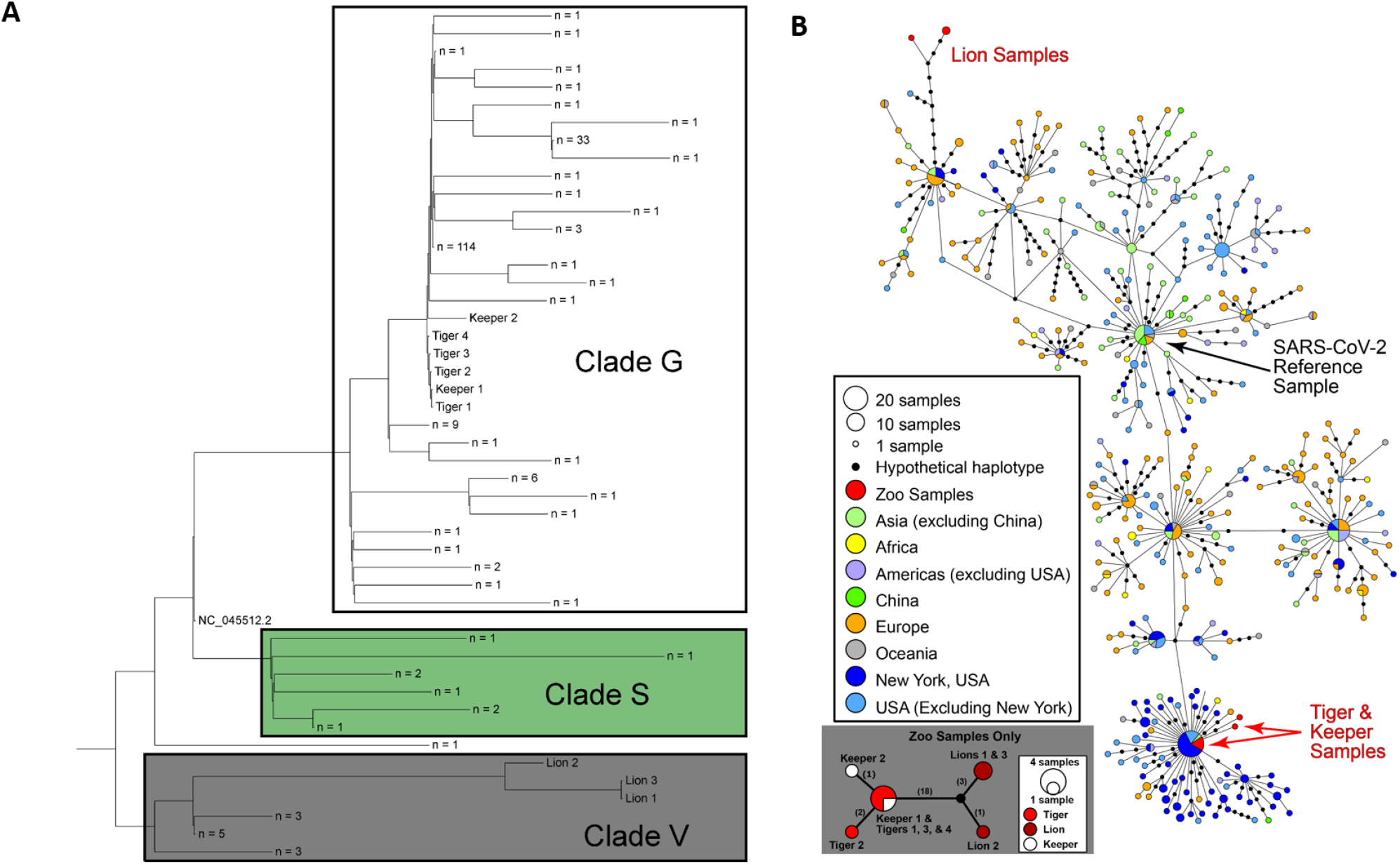
Phylogenetic and haplotype network analysis of SARS-CoV-2 strains obtained from tigers, lions and keepers. (A) Whole genome phylogeny of zoo SARS-CoV-2 sequences with Wuhan-Hu-1 reference genome (NC_045512-2) and consensus sequences of other publicly available sequences from New York clustered at 99.99% identity. The ML tree was unrooted then midpoint rooted. (B) Median-joining haplotype network analysis indicating relatedness and levels of genetic variation from a global dataset of 500 SARS-CoV-2 genomes. Differences are indicated by 1 step edges (lines) between black dots, which represent hypothetical or unsampled haplotypes. showing relatedness and levels of genetic variation from sequences obtained/generated from tigers, lions and keepers. Numbers in parentheses indicate differences between unique sequences (haplotypes). Black dot = hypothetical sequence not sampled.

The SARS-CoV-2 genome sequence from Tiger 1 was identical to the viral sequence recovered from Keeper 1 (a tiger keeper) and to other human SARS-CoV-2 strains detected in New York (NY_2929 [MT304486] and NY-QDX-00000001 [MT452574.1]) (**Fig. 3)**. These observations, temporal overlap in animal and human infections, and a lack of new animal introductions to the collection support the conclusion of transmission from an infected keeper(s) to the tigers. Whether this was direct or indirect (e.g. fomite, food handling/preparation) and whether subsequent tiger to tiger transmission (aerosol, respiratory droplet, etc.) occurred was not determined. A clear association and transmission source was not identified for the lions. The lion SARS-CoV-2 sequences were more divergent than those in the tigers and keepers (**Fig. 3**). However, nine of the 12 SNPs (relative to the Wuhan-Hu-1 reference strain) shared by all three lion viruses were also found in the closest human strain (GISAID accession: EPI_ISL_427161), which was detected nearby in Connecticut, US. Two lion keepers were serologically positive for SARS-CoV-2, but viral RNA was not detected and SARS-CoV-2 strain(s) could not be confirmed in their respiratory samples. However, given regular close contact between keepers and animals it is possible that the lions were also directly or indirectly infected by asymptomatic keepers or staff.

The host range of SARS-CoV-2 and other coronaviruses is determined primarily by the interaction of the virus S glycoprotein, specifically the spike 1 subunit (S1), and the cellular receptor, angiotensin-converting enzyme II (ACE2) (*12*). Recently, *in silico* predictions have shown high binding potential between the S receptor binding domain (RBD) and domestic cat ACE2 receptor, and that three of five amino acid residues that are critical for interaction with the SARS-CoV-2 S glycoprotein are conserved between human and domestic cat ACE2 (*12*). These observations are supported by reports describing natural and experimental infection of domestic cats with SARS-CoV-2 (*9*, *13*) and the data herein that shows a high degree of conservation between ACE2 in humans and domestic and wild felids (**Fig. S3**). Compared with the Wuhan-Hu-1 strain, the tiger and lion SARS-CoV-2 S gene sequences have 1-3 nucleotide and 0-2 nucleotide differences, respectively (**Fig. 4, Tables S9** and **S10**). These changes resulted in eight non-synonymous substitutions (P139L, L455S, F456Y, G496D, Q613R, D614G, A623T, and T716I). Notably, of three non-synonymous substitutions (L455S, F456Y, G496D) in the tiger strains, only one (G496D) was found in available human SARS-CoV-2 strains. These changes were not observed in the viral sequences from the lions (**Fig. 4**). Further work is needed to determine if these changes affect SARS-CoV-2 receptor binding and pathogenicity in felids and humans.

**Fig. 4.**
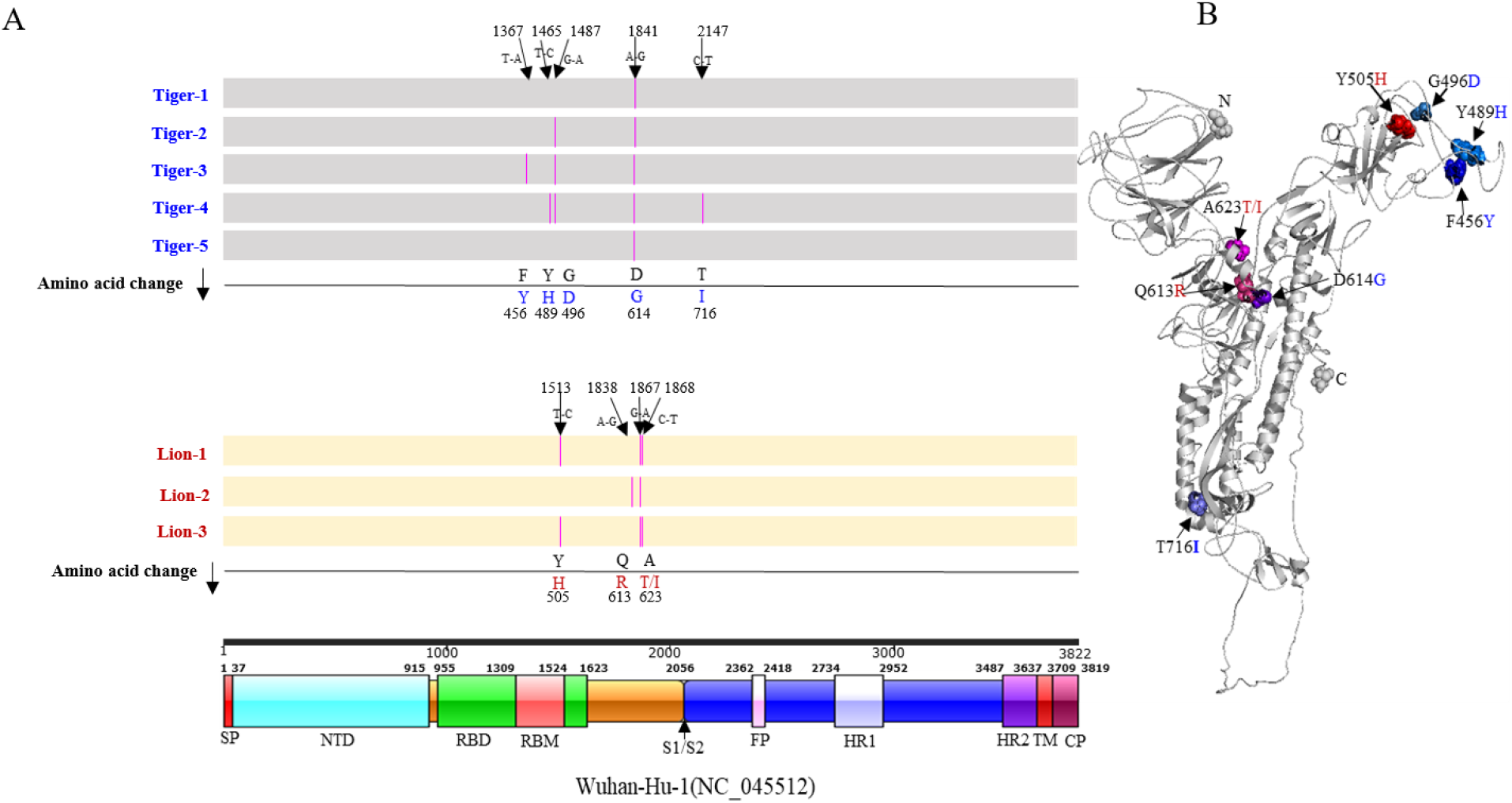
Nucleotide sequence and amino acid changes in the spike (S) protein of SARS-CoV-2 tigers and lions (A) Comparison with Wuhan-Hu-1 (NC_045512) (SnapGene v4.2.4). The nucleotide changes in each strain result in non-synonymous substitutions (pink lines) that are listed in line with the amino acid change. Schematic representation of the genome organization and functional domains of the S protein for SARS-CoV2 Wuhan-Hu-1 (IBS online software). (B) Structural modeling of the SARS-CoV-2 spike protein. Homology modeling of the Tiger 1 (MT365033) spike protein (I-TASSER [Iterative Threading ASSembly Refinement]). S protein amino acid changes in the tiger and lion strains versus the Wuhan-Hu-1 strain are depicted by pink lines from the N to C termini (F456Y, blue; Y489H, marine; G496D, sky blue; Y505H, red; Q613R, warm pink; D614G, purple/blue; A623T/I, magenta; T716I, slate).

Infections in the tigers and lions occurred at a time before testing was widely available in the US and there was limited evidence of pre-or asymptomatic viral shedding (*14*). They also preceded Centers for Disease Control and Prevention (CDC) guidance recommending face coverings (issued on April 3, 2020) to limit SARS-CoV-2 transmission. Keepers caring for the tigers and lions did not generally wear personal protective equipment (PPE) given the (historical) low risk of infectious respiratory disease transmission between humans and domestic or non-domestic felid species. Results of this investigation prompted the immediate development of new protocols for PPE use in the enclosures of non-domestic felids and other known or susceptible species including mustelids, viverrids, and chiroptera (PPE was already in place for work with non-human primates) at the Bronx Zoo. They also contributed to the development of similar recommendations by zoo and wildlife organizations (https://www.aza.org/aza-news-releases/posts/aza-and-aazv-statement-on-covid-19-positive-tiger-in-new-york).

The role of domestic and wild animal species in the epidemiology of SARS-CoV-2 is incompletely understood. To date, the reported number of cases of SARS-CoV-2 infection in domestic and wild animal species is low and, to our knowledge, no other zoos worldwide have confirmed cases in their animals. This is notable when considered in the context of the large number of human cases and close interactions between people, their pets and wild animals in their care. However, the fact that companion animals, farmed mink and zoo animals are susceptible to SARS-CoV-2 infection and shed infectious virus in respiratory secretions and/or feces (*7*, *9*) makes the human-animal interface an important area for further studies to elucidate transmission mechanisms and identify potential reservoirs of infection.

In the last two decades, at least three major coronavirus epidemics (SARS, MERS and COVID-19) have occurred. A feature shared by these and transmission of other novel viruses to humans including Ebola and HIV is an origin in a wild animal host. Despite a traditionally held perception of low risk, scientists and conservationists around the world have long recognized and shared concerns related to human activities that increase human/wildlife interactions and zoonotic disease transmission risk (*15*, *16*). As long as anthropogenic development and population growth bring humans and wildlife into ever increasing proximity, legal and illegal harvesting persists, and consumption of wildlife and wildlife products exists, there will be continued and significant risk of pandemic viral emergence with devastating global impact on human and animal health, economies, food security, and biodiversity.

## Supporting information

SARS-CoV-2 in tigers and lions

## Acknowledgements

The authors thank staff in the Bronx Zoo’s Clinical, Pathology, and Mammalogy Departments, and from Cornell AHDC Christopher Shiprack, Roopa Venugopalan and Renee Anderson for their help with sample collection and processing. They also wish to thank Dr. Eric Nelson at South Dakota State University and Beth Plocharczyk at Cayuga Medical Center for providing reagents used in the study. The following reagent was deposited by the Centers for Disease Control and Prevention and obtained through BEI Resources, NIAID, NIH: Genomic RNA from SARS-Related Coronavirus 2, Isolate USA-WA1/2020, NR-52285.

## Author contributions

Conceptualization, Project administration, Supervision: Denise McAloose, Leyi Wang, Suxiang Tong, Sally Slavinski, Paul P. Calle, Karen Terio, Mia Kim Torchetti, Diego G. Diel. Methodology, Validation, Formal Analysis, Investigation, Data Curation, Writing – Review & Editing: Melissa Laverack, Leyi Wang, Mary Lea Killian, Leonardo C. Caserta, Fangfeng Yuan, Patrick K. Mitchell, Krista Queen, Matthew R. Mauldin, Brittany D. Cronk, Susan L. Bartlett, John M. Sykes, Stephanie Zec, Tracy Stokol, Karen Ingerman, Martha A. Delaney, Marina Ivančić, Melinda Jenkins-Moore, Katie Mozingo, Kerrie Franzen, Nichole Hines Bergeson, Laura Goodman, Haibin Wang, Ying Fang, Colleen Olmstead, Colleen McCann, Patrick Thomas, Karen Terio, Mia Kim Torchetti, Diego G. Diel. Resources: Denise McAloose, Mary Lea Killian, Susan L. Bartlett, John M. Sykes, Stephanie Zec, Karen Ingerman, Ying Fang, Richard Frederikson, Erin Goodrich, François Elvinger, David C. Smith, Suxiang Tong, Sally Slavinski, Karen Terio, Paul P. Calle, Diego G. Diel. Writing – Original Draft Preparation, Review & Editing: Denise McAloose, Diego G. Diel. Funding Acquisition: Leyi Wang, Richard Frederikson, Laura Goodman, François Elvinger, Suxiang Tong, Paul P. Calle, Mia Kim Torchetti, Diego G. Diel.

## Data and materials availability

All data is available in the main text or the supplementary materials.

