## Supplementary material for "From people to *Panthera*: Natural SARS-CoV-2 infection in tigers and lions at the Bronx Zoo": SARS-CoV-2 in tigers and lions

**This PDF file includes:**

Materials and Methods

References 1 to 18

Figs. S1 to S3

Tables S1 to S9

Data S1 to S4

### Materials and Methods

#### Sample collection

Respiratory samples (nasal swab, oropharyngeal swab and tracheal wash) and blood were collected on April 2, 2020 from a female Malayan tiger (*Panthera tigris jacksoni*; Tiger 1) at the Wildlife Conservation Society's (WCS) Bronx, Zoo, New York, NY, United States (US). The tiger had developed an intermittent cough and wheezing on March 27, 2020. Physical examination, thoracic radiography and sample collection was performed with the tiger under general anesthesia. Additionally, voided fecal samples were collected opportunistically on April 4 and 5, 2020 from Tiger 1, all other tigers housed in the same building as Tiger 1 (Malayan tiger [Tiger 2], Amur tigers [*P. t. altaica*; Tigers 3, 4 and 5]), and three African lions (*P. leo krugeri* [Lions 1, 2 and 3]) housed in a different building. Tigers were housed individually but shifted between internal housing and outdoor exhibits through common spaces; lions were housed individually but were exhibited outdoors in alternating pairs in a shared environment. All except Tiger 5 developed respiratory signs similar to those presented by Tiger 1 in the week following disease onset in Tiger 1. Personal protective equipment (PPE), including N95 or surgical masks, face-shields or goggles, and disposable gloves, were worn during animal handling and sample collection. Diagnostic specimens were submitted to the University of Illinois Veterinary Diagnostic Laboratory (UIUC-VDL) and Animal Health Diagnostic Center at Cornell University (Cornell AHDC) (both part of the National Animal Health Laboratory Network) for broad diagnostic investigation (Tiger 1 - respiratory samples) and specific SARS-CoV-2 testing (all animals – fecal samples).

Epidemiologic investigation

Subsequent to the development of clinical signs and positive test results in the tigers and lions, an epidemiologic investigation into possible human infections in staff working with these animals was conducted by the New York City (NYC) and New York State (NYS) public health laboratories in conjunction with the US Centers for Disease Control and Prevention (CDC). The investigation focused on the time-period between March 16, 2020 (date on which the zoo was closed to the public due to the pandemic) and March 27 to April 1, 2020 (the timeline of disease onset in the animals). Twelve staff members (10 keepers, 2 managers) were identified that had responsibilities that offered opportunities for close (≤6 ft) but not direct contact with the animals during this time-period. This included moving animals between enclosures and exhibits, feeding, training sessions, enrichment activities, greetings and social interactions (e.g., chuffing - a form of vocalization that tigers performed that involves air exhalation, and which keepers also use to greet tigers). Additionally, lion keepers worked at a desk that was situated less than 6 feet from open fronted lion enclosures. Four keepers reported being mildly symptomatic (including fever, cough, chills, myalgia and fatigue) with signs beginning in each prior to or concurrently with illness in animals on March 20, 22, 27 and 28, 2020. Oropharyngeal swab and blood samples were collected on April 6, 2020 from the symptomatic keepers and SARS-CoV-2 specific rRT-PCR and a microsphere immunoassay (to detect IgG antibodies) were performed; sampling and testing were not performed in eight staff that did not report symptoms. All four of the tested keepers had evidence of SARS-CoV-2 infection (one rRT-PCR positive tiger keeper [Keeper 1], one rRT-PCR and serology positive tiger keeper [Tiger 2], two serology positive lion keepers [Keeper 3 and Keeper 4]). None of the keepers reported being sick at work and all stayed home while ill in compliance with organizational COVID-19 policies. rRT-PCR positive specimens were forwarded to CDC for whole genome sequencing (WGS) and haplotype network analysis to characterize the human samples and further compare the human and animal viral genome sequences. Interviews with the tiger and lion keepers suggested up to two additional keepers may have had signs or symptoms suggestive of mild and transient COVID-19; however, they did not self-report being sick and may not have recognized their symptoms as being consistent with COVID-19 and were not tested.

Non-SARS-CoV-2 feline respiratory pathogen testing

Nucleic acid extracted from the respiratory specimens (nasal and oropharyngeal swabs and tracheal wash fluid) from Tiger 1 were tested by qPCR or rRT-PCR for several common feline respiratory pathogens (Cornell AHDC) including *Bordetella bronchiseptica* (*1*), Influenza A virus (CDC universal assay (*2*), *Mycoplasma cynos* (*3*), *M. felis* (*4*), pneumovirus (*5*) with probe modification FAM-CTTCATCACTTTTGGCCTGGCCCAG-BHQ1), and S*treptococcus equi* supsp. *zooepidemicus* (*6*) **(Table S1)**. Additionally, samples were tested for *Chlamydia* spp. using a conventional PCR assay (*7*) **(Table S1)**. All assays have been adapted, optimized and are used routinely in feline infectious respiratory disease diagnostic testing.

Cytologic analysis of tracheal wash fluid

Direct smears of buoyant, flocculent material in the tracheal wash fluid and cytocentrifuge smears of the remaining fluid were prepared and stained with a Romanoski stain (modified Wright's-stain) using an automated stainer (Hema-tek 1000, Siemens). The stained slides were examined using standard brightfield microscopy by a board-certified veterinary clinical pathologist (Cornell AHDC).

*In situ* hybridization for SARS-CoV-2 RNA

Unstained cytologic smears of tracheal wash fluid from Tiger 1 were fixed in ice-cold 100% methanol for 20 minutes and stored at -80°C until shipping to UIUC College of Veterinary Medicine Zoological Pathology Program (ZPP) on cold packs. Upon arrival, the slides were submerged for an additional 30 minutes in 10% neutral buffered formalin at 4 ºC, air-dried, then placed in 100% ethyl alcohol for 5 minutes at room temperature. An *in situ* hybridization (ISH) chromogenic manual assay was performed using the RNAScope® 2.5 HD Detection Kit Red and a 20-pair oligonucleotide probe targeted to the SARS-CoV-2 S gene of the Wuhan Hu-1 complete genome (NC_045512.2; Advanced Cell Diagnostics # 848561**)** according to the manufacturer's directions (Advanced Cell Diagnostics, Inc., Newark, CA). Positive control slides consisted of Vero cells infected with SARS-CoV-2 isolated from Tiger 1. A control probe targeting the DapB gene from the *Bacillus subtilis* strain SMY (Advanced Cell Diagnostics # 310043) was used as negative control on all cytology sections in parallel with the SARS-CoV-2 target probe **(Fig. S1)**. Additional negative controls to rule out cross-reactivity with tiger RNA and the felid alphacoronavirus included: a cytocentrifuge preparation of cell cultures infected with the alphacoronavirus feline enteric coronavirus (FeCoV) (kindly provided by Dr. Gary Whittaker, Cornell University); formalin-fixed, paraffin-embedded (FFPE) unstained sections of normal Malayan tiger trachea, lung and oropharyngeal tissue; and lung, lymph node and intestine from a FeCoV positive domestic cat **(Fig. S1)**. Samples from the control tiger and domestic cat were collected opportunistically in 2014 and 2013, respectively, and archived as part of routine necropsy procedures (Wildlife Conservation Society's (WCS) Bronx Zoo and UIUC-ZPP, , respectively).

Virus isolation

Virus isolation on respiratory and fecal samples was performed in Vero (ATCC CCL-81), Vero E6, and Vero 76 cells under Biosafety Level-3 conditions at the Cornell AHDC and the National Veterinary Services Laboratory (NVSL). Cells were cultured in minimum essential medium eagle (MEM-E; Gibco, Gaithersburg, MD) supplemented with 2.5-10% fetal bovine serum (FBS; Gibco), 100 IU/mL penicillin, and 100 µg/mL streptomycin (growth media). Cells were seeded in 12-well culture plates or T25 flasks and cultured at 37$^o$C with 5% $CO_2$ for 24 - 48h. Before inoculation, respiratory swabs and tracheal wash fluid samples were diluted at 1:10, 1:100 and 1:1000 in serum-free MEM-E containing 200 UI/mL, penicillin, 200 µg/mL streptomycin, and 2.5 µg/mL Amphotericin B (all from Gibco); swabs from fecal samples were placed in 1 ml of sterile PBS supplemented with 2.5% BSA (Sigma Aldrich, St Louis, MO) containing 200 UI/mL, penicillin, 200 µg/mL streptomycin and 2.5 µg/mL Amphotericin B (all from Gibco). Cells were rinsed with MEM-E and inoculated with 300 µl of each respiratory sample dilution in individual wells of a 12-well plate or 1.5 ml of the diluted fecal sample in a T25 flask and adsorbed for 1 h 37$^o$C with 5% $CO_2$ for 1 h. Mock-inoculated cells were used as negative controls. After adsorption, replacement medium was added and cells were incubated at 37$^o$C with 5% $CO_2$ and monitored daily for cytopathic effect (CPE) for five days. Cell cultures with no CPE were frozen, thawed, and subjected to three blind passages with inoculation of fresh Vero cell cultures with the lysates as described above. SARS-CoV-2 infection in CPE positive cultures was confirmed with SARS-CoV-2-specific rRT-PCR using the CDC N1 primer and probe set, an immunofluorescence assay using a mouse monoclonal antibody against the SARS-CoV N protein (*8*, *9*), and RNAscope® *in situ* hybridization as described above.

Virus neutralization (VN) assay

Seroconversion of Tiger 1 to SARS-CoV-2 was assessed by a virus neutralization assay (VN; Cornell AHDC). Two-fold serial dilutions (1:8 to 1:4096) of a serum sample collected on April 2, 2020 (6 days after the onset of clinical signs) were incubated with 100 tissue culture infective dose ($TCID_{50}$) of SARS-CoV-2 for 1 h at 37°C. Following incubation of serum and virus, 50 µl of a cell suspension of Vero CCL-81 cells was added to each well of a 96-well plate and incubated for 72 h at 37°C with 5% $CO_2$. Neutralizing antibody titers were expressed as the reciprocal of the highest dilution of serum that completely inhibited CPE. Archived frozen sera from another tiger (Cornell AHDC) and positive human control sera (de-identified convalescent human sera provided by Cayuga Medical Center, IRB protocol 0420EP) were included and all samples were tested in triplicate with results averaged. A cell culture control was included in the assays and the virus working dilution was back-titrated.

SARS-CoV-2 real-time reverse transcriptase PCR (rRT-PCR)

Nucleic acid was extracted from nasal and oropharyngeal swabs and tracheal wash fluid (Tiger 1) using the MagMAX isolation kit (Thermo Fisher Scientific, Waltham, MA) and from fecal samples (Tigers 1-5 and Lions 1-3) using the MagMax Core nucleic acid purification kit (Thermo Fisher Scientific, Waltham, MA) and an automated nucleic acid extractor (King Fisher Flex Purification System, Thermo Fisher Scientific, Waltham, MA) according to the manufacturer's instructions. Analysis for SARS-CoV-2 RNA was performed with rRT-PCR and either the 2019-nCoV CDC qPCR Probe Assay targeting three regions of the nucleocapsid (N) gene (N1, N2, and/or N3; Integrated DNA Technologies [IDT], Inc., Coralville, IA) (UIUC-VDL, Cornell AHDC) or the CDC qPCR Probe Assay and in-house primers for the envelope (E) gene with the AgPath-ID One-Step RT-PCR Kit (Applied Biosystems, Foster City, CA) (UIUC-VDL). Primers and probe sequences are listed in **Table S3**. Thermal cycler conditions consisted of reverse transcription and enzyme activation at 45 - 48°C for 10 min and 95°C for 10 min, respectively, followed by 40 - 45 cycles of 95°C for 3 - 15 sec and 55 - 60°C for 45 sec. Positive (2019-nCoV_N_Positive Control, IDT Coralville, IA and a synthesized plasmid [GenScript] E gene control) and negative controls (distilled or nuclease-free water), plus internal amplification controls (Xeno or beta actin; Thermo Fisher Scientific, Waltham, MA) were included as separate reactions. Following initial rRT-PCR testing at both institutions, samples were submitted to the NVSL for confirmatory testing, using the 2019-nCoV CDC qPCR Probe Assay and N1 and N2 primers.

Amplicon sequencing

MinION and Sanger amplicon sequencing was used to confirm rRT-PCR results at Cornell AHDC and NVSL, respectively. Primer sequences are listed in **Table S3.** For MinION-based sequencing, targets were amplified directly from tracheal wash or fecal samples using the SuperScript™ IV One-Step RT-PCR System (Thermo Fisher Scientific, Waltham, MA). Primers targeted the complete spike (S) gene (4023 bp) and an internal region of the N gene (634 bp). Universal Oxford nanopore-compatible adapter sequences were added to the 5' end of each primer sequence to allow PCR-based barcoding. Amplicons were purified (AMPure XP beads [Beckman Coulter, Brea, CA]; 1.6:1 volumetric bead-to-DNA ratio) and DNA quantification was performed on a Qubit® fluorometer 3.0 (dsDNA High Sensitivity Assay kit; Thermo Fisher Scientific, Waltham, MA). Samples were subsequently diluted to 0.5 nM in a total of 24 µl and used as the input for the library preparation following the 1D PCR barcoding (96) genomic DNA

(SQK-LSK109) protocol (Oxford Nanopore Technologies, Oxford, UK). Final DNA libraries were loaded in a FLO-MIN106 R9.4 flow cell to start the sequencing runs.

Sanger sequencing was performed using primers targeting partial regions of S, N and RNA dependent RNA polymerase (RdRp) genes. Amplicons were generated directly from nasal and oropharyngeal swabs and tracheal wash fluid using the SuperScript™ III One-Step RT-PCR System (Thermo Fisher Scientific, Waltham, MA). Reactions were purified using the Qiagen PCR Purification Kit (Qiagen, Germantown, MD). DNA was amplified for Sanger sequencing using the BigDye Terminator v 3.1 Cycle Sequencing Kit (Thermo Fisher Scientific, Waltham, MA), and sequenced on the Applied Biosystems 3500xl Genetic Analyzer (Thermo Fisher Scientific, Waltham, MA).

Whole Genome Sequencing and phylogenetic analysis

Whole genome sequencing (WGS) was performed on tracheal wash fluid and fecal specimens from all individual tigers and lions as previously described (*10*). Individual fecal samples from Tigers 1-5 and Lions 1-3 and cell culture viral isolates were subjected to sequencing with either MinION-based amplicon sequencing using overlapping primers covering the full viral genome (amplicons with an average size of ~1,500 bp; primer sequences are listed in **Table S9**) or the Ion AmpliSeq Kit for Chef DL8 and Ion AmpliSeq SARS-CoV-2 Research Panel (Thermo Fisher Scientific, Waltham, MA). MinION libraries were prepared as previously described (*11*) using the Native Barcode Kit, EXP-NBD104, Ligation Sequencing Kit, SQK-SQK109 (Oxford Nanopore Technologies) and sequenced on a R9.4 flow cell for 6 hours. Ion targeted libraries were sequenced using an Ion 530 chip on the Ion S5 system using the Ion 510™ & Ion 520™ & Ion 530™ Kit (Thermo Fisher Scientific, Waltham, MA). Viral isolates were sequenced with the MinION or Ion AmpliSeq approach as above (**Data S3**). Whole genome sequencing on the rRT-PCR positive specimens from the two SARS CoV 2 positive keepers was performed as previously described using an amplicon sequencing approach and Sanger sequencing (*12*).

All genomes for each animal that were assembled using data generated from different sequencing platforms (Illumina, MinION and/or Ion Torrent) were combined into a single consensus sequence for each animal (**Data S2**). The assemblies for a given animal were aligned with the Wuhan-Hu-1 reference sequence (NC_045512.2) using MAFFT v. 7.453 (*13*). The reference sequence was then removed from the alignment and a consensus sequence for the virus sequence recovered from each animal was generated using the consambig program in EMBOSS v. 6.6.0.0 (*14*). When a single assembly shifted alignment due to a single base insertion or repeat nucleotide, the alignment was re-run after removal of the offending nucleotides.

To compare the outbreak genomes to others isolated from humans in the same geographic region, all available SARS-CoV-2 genomes from New York were downloaded from NCBI on April 23, 2020. These were clustered at 99.99% identity using vsearch v. 2.14.2 (*15*) and the consensus sequences from each cluster were aligned along with Wuhan-Hu-1 reference sequence, using MAFFT v. 7.453 (*13*). A phylogenetic tree was constructed using the consensus sequence from each animal, keepers, sequences from New York, and the Wuhan-Hu-1 reference sequence (NC_045512.2) using the GTR-Gamma model in RAxML v. 8.2.12 (*16*).

<u>Haplotype network analysis</u>

Haplotype network analyses were conducted with two overlapping datasets using PopART software (*17*) using the median joining algorithm (*18*). Dataset 1 contained nine genomes generated from the Bronx Zoo cases: four tigers, three lions, and two tiger keepers. Dataset 2 contained the nine genomes from Dataset 1, as well as 500 additional genomes (including the SARS-CoV-2 reference Wuhan-Hu genome) generated from a total of 53 countries to better understand genetic relatedness of Bronx Zoo cases in the context of the global pandemic. The top 10 BLAST results for the lion sequences were included in Dataset 2.

For both datasets, the entire genome alignment was examined visually for accuracy and evidence of large-scale rearrangements to rule out the likelihood of multiple single nucleotide polymorphisms (SNPs) being the result of a single evolutionary event. Subsequently, whole genome alignments were converted into a SNP matrix by removing columns containing identical bases, gaps, and ambiguous bases. Length of final SNP matrices were 25nt (Dataset 1) and 567nt (Dataset 2).

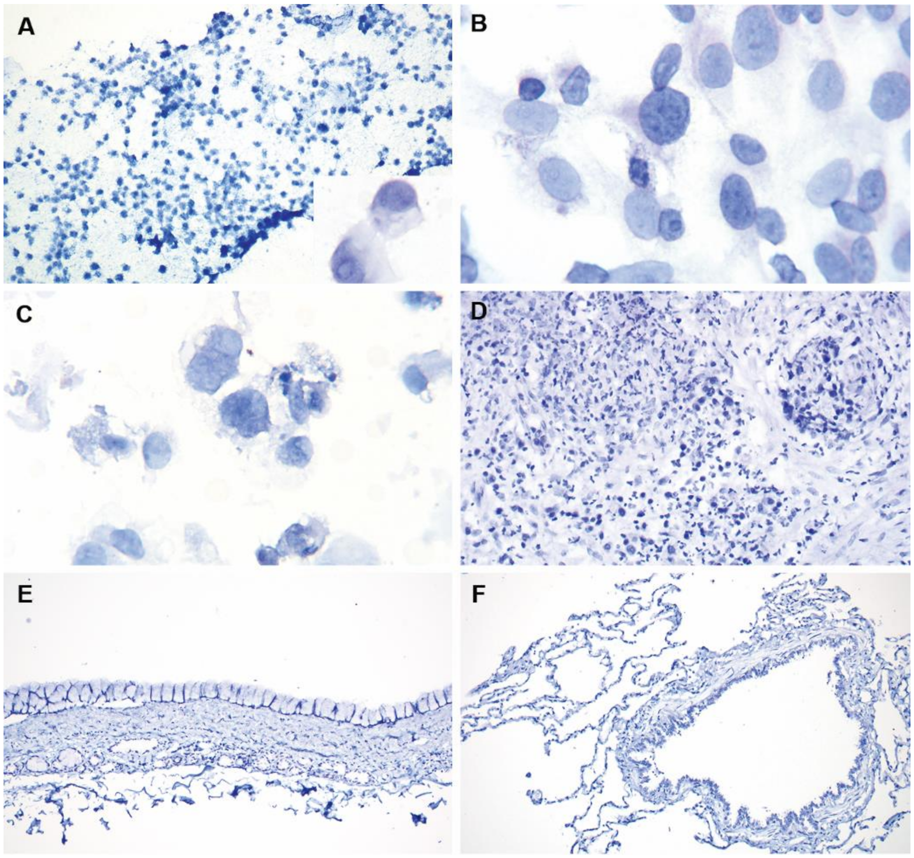


**Figure S1**. Negative controls for *in situ* hybridization (RNAScope® red chromogenic manual assay and hematoxylin counterstain). No specific staining is noted in any of the control sections confirming specificity of the probes for SARS-CoV-2 and lack of cross-reaction with normal tiger cells and feline alphacoronaviruses. (A, B) Direct smears of Tracheal wash from Tiger 1 (A) and infected Vero Cells (B) incubated with negative control probe targeting the DapB gene from the *Bacillus subtilis* strain SMY. (C) Cells infected with feline enteric coronavirus incubated with the probe targeting SARS-CoV-2. (D) Histologic section of mesentery from a domestic cat with confirmed feline infectious peritonitis (the mutated form of FeCoV) incubated with the probe targeting SARS-CoV-2. (E, F) Histologic sections of normal trachea (E) and lung

(F) from a Malayan tiger collected prior to the emergence of SARS-CoV-2 incubated with the probe targeting SARS-CoV-2.

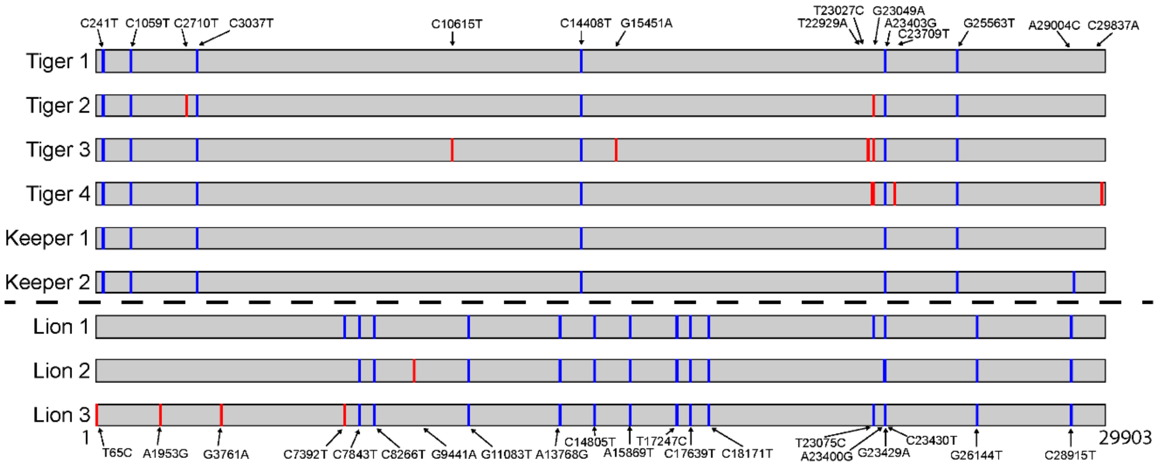


**Figure S2.** SNP sites relative to Wuhan-Hu-1 genome. SNPs for tiger and keeper sequences are labeled along the top and SNPs for lion sequences are labeled along the bottom. Blue bars indicate a unanimous consensus nucleotide across all assemblies for a given animal or keeper and red bars indicate ambiguous base calls within that individual.

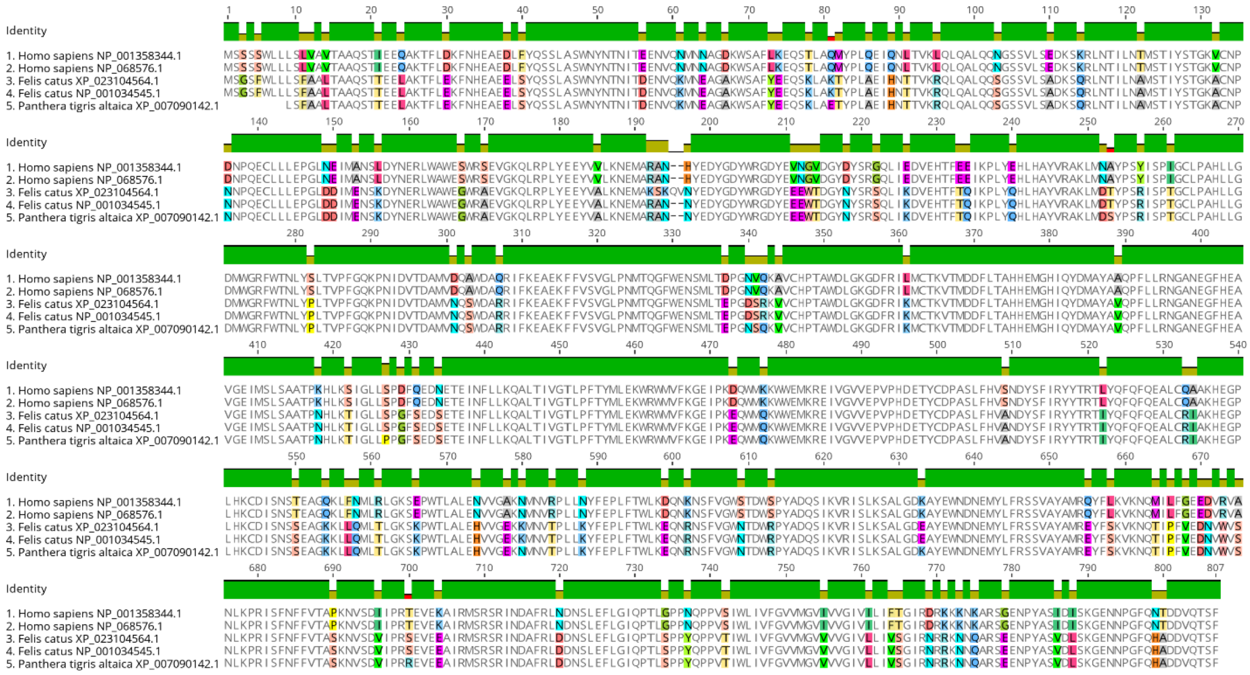


**Figure S3.** Alignment of human, domestic cat and tiger in ACE2 amino acid sequences. Amino acid alignment was performed using MAFFT (*13*) with parameters Auto for algorithm, BLOSUM62 for scoring matrix, Gap open penalty of 1.53 and Offset value of 0.123. Positions where amino acid differences were noted are highlighted in colors.

**Table S1.** PCR results for common feline respiratory pathogens in tracheal wash fluid, nasal (NS) and oropharyngeal (OP) swabs from Tiger 1 in a standard feline diagnostic panel performed by the Cornell AHDC.

| PCR target | Specimen | Result | Specimen | Result |
|---|---|---|---|---|
| Influenza A virus | Tracheal wash | Not detected | NS/OP | Not detected |
| Pneumovirus | Tracheal wash | Not detected | NS/OP | Not detected |
| *Mycoplasma cynos* | Tracheal wash | Not detected | NS/OP | Not detected |
| *Mycoplasma felis* | Tracheal wash | Not detected | NS/OP | Not detected |
| *Bordetella* spp. | Tracheal wash | Not detected | NS/OP | Not detected |
| *Streptococcus zooepidemicus* | Tracheal wash | Not detected | NS/OP | Not detected |
| *Chlamydia* spp.[a] | Tracheal wash | Not detected | NS/OP | Not detected |
| **Virus isolation[b]** | **Specimen** | **Result** | **Specimen** | **Result** |
| Feline herpesvirus | Tracheal wash | No virus isolated | NS/OP | No virus isolated |
| Feline calicivirus | Tracheal wash | No virus isolated | NS/OP | No virus isolated |

[a]This PCR detects *C. psittaci*, *C. felis* and *C. abortus*. [b]Virus isolation was performed in feline lung cells as part of the routine feline respiratory panel screening at Cornell AHDC.

**Table S2.** Primers and probes used for rRT-PCR detection and MinION or Sanger amplicon sequencing of SARS-CoV-2 from tiger nasal, oropharyngeal, tracheal wash and fecal samples and lion fecal samples at three independent laboratories.

| Target | Primers/Probes | Amplicon Size | Application | Sequence | Laboratory |
|---|---|---|---|---|---|
| Nucleocapsid | 2019-nCoV_N1-F | n/a | rRT-PCR | 5'-GACCCCAAAATCAGCGAAAT-3' | Cornell AHDC, NVSL |
| | 2019-nCoV_N1-R | | | 5'-TCTGGTTACTGCCAGTTGAATCTG-3' | |
| | 2019-nCoV_N1-P | | | 5'-FAM-ACCCCGCATTACGTTTGGTGG ACC-BHQ1-3' | |
| Nucleocapsid | 2019-nCoV_N2-F | | rRT-PCR | 5'-TTACAAACATTGGCCGCAAA-3' | Cornell AHDC, UIUC-VDL, NVSL |
| | 2019-nCoV_N2-R | | | 5'-GCGCGACATTCCGAAGAA-3' | |

| | | | | | |
|---|---|---|---|---|---|
| | 2019-nCoV_N2-P | | | 5’-FAM-ACAATTTGCCCCCAGCGCTTC AG-BHQ1-3’ | |
| Nucleocapsid | 2019-nCoV_N3-F | | rRT-PCR | 5’-GGGAGCCTTGAATACACCAAAA-3’ | Cornell AHDC |
| | 2019-nCoV_N3-R | | | 5’-TGTAGCACGATTGCAGCATTG-3’ | |
| | 2019-nCoV_N3-P | | | 5’-FAM-AYCACATTGGCACCCGCAATC CTG-BHQ1-3’ | |
| Envelope | E-F | | rRT-PCR | 5’-ATGGAACCAATTTATGATGAACCG-3’ | UIUC-VDL |
| | E-R | | | 5’-CGAATGAGTACATAAGTTCGTAC-3’ | |
| | E-P | | | 5’-56-FAM/ACGACGACTACTAGCGTG CC/36-tamsP-3’ | |
| Spike | SARS-CoV S-F | 4023 | MinION | 5’-TTTCTGTTGGTGCTGATATTGCAG GGGTACTGCTGTTATGTCT-3’ | Cornell AHDC |
| | SARS-CoV-S-R | | | 5’-ACTTGCCTGTCGCTCTATCTTCGC GCGAACAAAATCTGAAGG-3’ | |

| | | | | | |
|---|---|---|---|---|---|
| Nucleocapsid | SARS-CoV N int-F | 634 | MinION | 5’-TTTCTGTTGGTGCTGATATTGCATCAC ATTGGCACCCGCAAT-3’ | Cornell AHDC |
| | SARS-CoV N int-R | | | 5’-ACTTGCCTGTCGCTCTATCTTCAGGCT CTGTTGGTGGGAATG-3’ | |
| Spike | Spike-F | 443 | Sanger | 5’-GAAGACCCAGTCCCTACTTATTG-3’ | NVSL |
| | Spike-R | | | 5’-GCTGTCCAACCTGAAGAAGA-3’ | NVSL |
| Nucleocapsid | 2019-nCoV_N1-F | 466 | Sanger | 5’-GACCCCAAAATCAGCGAAAT-3’ | NVSL |
| Nucleocapsid | 2019-cCoV_N3-R | | | 5’-TGTAGCACGATTGCAGCATTG-3’ | |
| RdRp | RdRp_SARSr-F | 410 | Sanger | 5’-GTGARATGGTCATGTGTGGCGG-3’ | NVSL |
| RdRp | COVID-410-R | | | 5’-CCAACATTTTGCTTCAGACATAAAAAC-3’ | |

N = nucleocapsid gene; N int = internal region of the N gene; E = envelope gene; S = spike protein gene; RdRp = RNA dependent RNA polymerase; F = forward; R = reverse; P = probe; int: internal primers; Underlined sequenced in MinION primers represent adapter sequences added to each primer.

**Table S3.** Average Ct values from real-time RT-PCR results for nasal and oropharyngeal swabs and tracheal wash fluid from Tiger 1 performed at three independent laboratories using primers targeting the nucleocapsid gene (N1, N2, N3) using the 2019-nCoV CDC qPCR Probe Assay and in-house primers for the envelope (E) protein.

| | Cornell AHDC | | | UIUC_VDL | | NVSL | |
|---|---|---|---|---|---|---|---|
| Sample/Target | N1 | N2 | N3 | N2 | E | N1 | N2 |
| Nasal | 21.26[a] | 20.99 | 20.59 | 19.94 | 28.13 | 18.20 | 18.30 |
| Oropharyngeal | | | | 28.23 | 35.00 | 27.00 | 27.60 |
| Tracheal wash 1 | 20.97[b] | 21.00 | 20.46 | 21.49 | 29.01 | 20.05 | 20.35 |
| Tracheal wash 2 | | | | 21.73 | 28.71 | 20.25 | 20.10 |

[a]Pooled nasal and oropharyngeal swab samples were tested.

[b]Pooled aliquots from a single tracheal wash fluid sample were tested.

UIUC-VDL, Cornell AHDC Ct cutoff values: N1, N2, N3 < 40; E < 37

NVSL, all samples were run in duplicate. Ct cutoff values: N1, N2 < 40.

**Table S4.** Average Ct values from real-time RT-PCR results for fecal samples from five tigers and two lions collected on day eight and nine after Tiger 1 first demonstrated respiratory signs. The RT-PCR was performed at two institutions using primers targeting the nucleocapsid (N1, N2, N3) gene with the 2019-nCoV CDC qPCR Probe Assay and in-house primers for the envelope (E) gene.

| Laboratory | Cornell AHDC | Cornell AHDC | Cornell AHDC | UIUC-VDL | UIUC-VDL |
|---|---|---|---|---|---|
| Sample Collection Date | 4-Apr-20 | 4-Apr-20 | 4-Apr-20 | 5-Apr-20 | 5-Apr-20 |
| Animal ID/rRT-PCR | N1 | N2 | N3 | N2 | E |
| Tiger 1 | NA | NA | NA | 23.32 | 33.54 |
| Tiger 2 | 22.28 | 25.83 | 24.18 | 14.48 | 26.08 |
| Tiger 3 | 24.56 | 29.51 | 26.18 | 27.37 | 34.00 |
| Tiger 4 | 27.91 | 31.72 | 29.25 | 24.67 | 35.22 |
| Tiger 5 | 32.03 | 36.28 | 32.87 | 28.84 | 0 |
| Lion 1 | 31.04 | 35.69 | 33.51 | 25.13 | 33.64 |
| Lion 2 | 23.19 | 26.17 | 23.84 | 17.29 | 28.63 |
| Lion 3 | 31.22 | 34.83 | 34.10 | 24.97 | 35.00 |

NA = sample not available.

Ct cutoff values: N1, N2, N3 < 40; E < 37

**Table S5.** Targeted MinION amplicon sequencing for the full SARS-CoV-2 spike (S) gene and an internal region of the nucleocapsid (N) gene to confirm the presence of viral RNA in tracheal wash fluid of Tiger 1 and fecal samples of Tigers 2-5 and Lions 1-3.

| Full S gene | | | | | |
|---|---|---|---|---|---|
| | N. of reads[a] | Mean depth of coverage | Minutes of sequencing[b] | BLAST search | Percent id. |
| Tiger 1[c] | 200 | 141.3 | 1 | LC547533.1 | 100 |
| Tiger 2[d] | 1066 | 1052.9 | 30 | LC547533.1 | 99.98 |
| Tiger 3[d] | 2405 | 2478.3 | 30 | LC547533.1 | 99.98 |
| Tiger 4[d] | 2921 | 2892.5 | 30 | LC547533.1 | 99.95 |
| Tiger 5[d] | 44 | 22.1 | 150 | LC547533.1 | 100 |
| Lion 1[d] | 30 | 14.8 | 150 | MT447170.1 | 100 |
| Lion 2[d] | 2432 | 2389.8 | 30 | MT447170.1 | 99.95 |
| Lion 3[d] | 77 | 27.2 | 150 | LC547533.1 | 100 |

| Partial N gene | | | | | |
|---|---|---|---|---|---|
| | N. of reads | Mean depth of coverage | Minutes of sequencing | BLAST search | Percent id. |
| Tiger 1[c] | 482 | 458.8 | 1 | LC547522.1 | 100 |
| Tiger 2[d] | 16323 | 16126.7 | 30 | LC547522.1 | 100 |
| Tiger 3[d] | 23455 | 23255.8 | 30 | LC547522.1 | 100 |
| Tiger 4[d] | 20141 | 19934.9 | 30 | LC547522.1 | 100 |

| | | | | | |
|---|---|---|---|---|---|
| Tiger 5[d] | 1479 | 1459.7 | 30 | LC547522.1 | 100 |
| Lion 1[d] | 109 | 106.8 | 30 | MT358693.1 | 99.84 |
| Lion 2[d] | 17513 | 17360.3 | 30 | MT358693.1 | 100 |
| Lion 3[d] | 1290 | 1273 | 30 | MT358693.1 | 100 |

[a] Number of reads mapped to genome used as reference (MN985325.1) after processing raw reads.

[b] Minutes necessary after starting sequencing run to generate consensus sequence.

[c]Tracheal wash fluid sample.

[d]Fecal sample.

**Table S6.** Virus isolation for SARS-CoV-2 and rRT-PCR confirmation in respiratory and fecal samples from tigers and lions.

| Laboratory | | Cornell AHDC | | | NVSL | | | Cornell AHDC | | |
|---|---|---|---|---|---|---|---|---|---|---|
| Sample Collection Date | | 4-Apr-20 | | | 4-Apr-20 | | | 8-Apr-20 | | |
| Animal ID/Passage (P) | Sample | P 1 | P 2 | P 3 | P 1 | P 2 | P 3 | P 1 | P 2 | P 3 |
| Tiger 1 | Tracheal wash | - | + | + | - | - | - | NA | NA | NA |
| Tiger 1 | Nasal/oropharyngeal swab | - | - | - | - | - | - | NA | NA | NA |
| Tiger 2 | Feces | - | - | - | - | - | - | - | - | - |
| Tiger 3 | Feces | - | - | - | - | - | - | - | - | + |
| Tiger 4 | Feces | - | - | - | - | - | - | - | - | - |
| Tiger 5 | Feces | - | - | - | - | - | - | - | - | - |
| Lion 1 | Feces | - | - | - | - | - | - | - | - | - |
| Lion 2 | Feces | - | - | - | - | + | + | - | - | - |
| Lion 3 | Feces | - | - | - | - | - | - | - | - | - |

NA = sample not available.

-: No virus isolated; +: SARS-CoV-2 isolated, with confirmation by rRT-PCR (all positive samples), immunofluorescence staining for the N protein, and *in situ* hybridization (Tiger 1, tracheal wash)

**Table S7.** Virus neutralization assay performed on serum taken from Tiger 1 collected on April 2, 2020, six days after respiratory signs were noted. Serial dilutions of the serum wereincubated with 100 $TCID_{50}$ of SARS-CoV-2 isolate TGR/NY/20 on Vero cells. Virus cytopathic effect was used as an indicator of virus infection/replication.

| Sample | 1:4 | 1:8 | 1:16 | 1:32 | 1:64 | 1:128 | 1:256 | 1:512 | 1:1028 | 1:2048 |
|---|---|---|---|---|---|---|---|---|---|---|
| Negative control[a] | + | + | + | + | + | + | + | + | + | + |
| Positive control[b] | - | - | - | - | - | - | + | + | + | + |
| Tiger 1 | - | - | - | - | - | + | + | + | + | + |

-: No CPE (inhibition of infection/replication); +: Presence of CPE. Antibody titers were defined as the reciprocal of the highest serum dilution that completely inhibited CPE (1:64 for Tiger 1).

[a] Serum from another tiger collected in 2019 and available in Cornell AHDC frozen archives.

[b] Convalescent human serum (IRB #0420EP).

**Table S8.** Comparison of nucleotide and amino acid mutations between viral strains sequenced from five tigers, three lions, two keepers (Keeper 1 and 2) and the Wuhan-Hu-1 strain across the entire SARS-CoV-2 genome.

| Position | Subject | Nucleotide Change* | Amino Acid Change** | Gene |
|---|---|---|---|---|
| 241 | All | C / T | NA | 5'UTR |
| 1059 | All | C / T | T / I | Nsp2 |
| 2710 | Tiger 2 | C / Y | No change | Nsp2 |
| 3037 | All | C / T | No change | Nsp3 |
| 10615 | Tiger 3 | C / Y | No change | 3C-like proteinase |
| 14408 | All | C / T | P / L | Nsp12 (RdRp) |
| 15451 | Tiger 3 | G / R | G / S | Nsp12 (RdRp) |
| 22929 | Tiger 3 | T / W | F / Y | S |
| 23027 | Tiger 4 | T / Y | Y / H | S |
| 23049 | Tiger 2<br>Tiger 3<br>Tiger 4 | G / R | G / D | S |
| 23403 | All | A / G | D / G | S |
| 23709 | Tiger 4 | C / Y | T / I | S |
| 25563 | All | G / T | Q / H | ORF3a |
| 29004 | Keeper 2 | A / C | Q / P | N |
| 29837 | Tiger 4 | C / M | NA | 3'UTR |

* For ambiguous base calls, the following ambiguity codes were used: Y = C or T; R = A or G;

M = A or C

** For ambiguous sites, amino acid change given for the non-reference base

**Table S9.** Primers used for target-based whole genome sequencing of SARS-CoV-2 strains from tigers and lions using MinION

| Primer name | Primer sequence |
|---|---|
| nCoV 29,860 R pool 1 | GGCTCTTCCATATAGGCAGCT |
| nCoV 28,362 R pool 2 | TTCGCTGATTTTGGGGTCCA |
| nCoV 28,235 F pool 1 | TGGGTAGTCTTGTAGTGCGT |
| nCoV 26,862 R pool 1 | GCTGAGCCACATCAAGCCTA |
| nCoV 26,736 F pool 2 | TTTCCTCTGGCTGTTATGGC |
| nCoV 25,290 R pool 2 | AGCCAGCTATAAAACCTAGCCA |
| nCoV 25,175 F pool 1 | CCTCAATGAGGTTGCCAAGA |
| nCoV 23,724 R pool 1 | CTGCACCAAGTGACATAGTGT |
| nCoV 23,573 F pool 2 | AGGGGCTGAACATGTCAACA |
| nCoV 22,274 R pool 2 | CTGAGGGAGATCACGCACTA |
| nCoV 22,151 F pool 1 | GGACCTTGAAGGAAAACAGGGT |
| nCoV 20,688 R pool 1 | TGGCCATCTTTACACCAAAGC |
| nCoV 20,518 F pool 2 | AACAGATGCGCAAACAGGTTC |
| nCoV 19,172 R pool 2 | TGTCACTACAAGGCTGTGCA |
| nCoV 19,015 F pool 1 | TGCGGCTTGTAGAAAGGTTCA |
| nCoV 17,640 R pool 1 | ACAATTTCAGCAGGACAACGC |
| nCoV 17,493 F pool 2 | GCGACCCTGCTCAATTACCT |
| nCoV 16,267 R pool 2 | AGCCTCATAAAACTCAGGTTCCC |
| nCoV 16,093 F pool 1 | TGCTTACCCACTTACTAAACATCCT |

| | |
|---|---|
| nCoV 14,844 R pool 1 | GCAGCATTACCATCCTGAGC |
| nCoV 14,636 F pool 2 | ATGCACGCTGCTTCTGGTAA |
| nCoV 13,442 R pool 2 | GCAGACGGTACAGACTGTGT |
| nCoV 13,280 F pool 1 | ATCCTTTGGTGGTGCATCGT |
| nCoV 12,002 R pool 1 | CTGGACACATTGAGCCCACA |
| nCoV 11,855 F pool 2 | GTTGGGTGTTGGTGGCAAAC |
| nCoV 10,508 R pool 2 | GGGCCTCATAGCACATTGGT |
| nCoV 10,403 F pool 1 | GACACCTAAGTATAAGTTTGTTCGC |
| nCoV 9,058 R pool 1 | GCTGATGTTGCAAAGTCAGTGT |
| nCoV 8,912 F pool 2 | TTTTGTCGTGCCTGGTTTGC |
| nCoV 7,592 R pool 2 | ACATTCGACTCTTGTTGCTCT |
| nCoV 7,446 F pool 1 | GCCCCGATTTCAGCTATGGT |
| nCoV 6,235 R pool 1 | TCAATAGCCACCACATCACCA |
| nCoV 6,082 F pool 2 | ATCCAAACGCAAGCTTCGAT |
| nCoV 4,716 R pool 2 | ACCGAGCAGCTTCTTCCAAA |
| nCoV 4,563 F pool 1 | GGTGTGGTTGATTATGGTGCT |
| nCoV 3,223 R pool 1 | GCAGAAGTGGCACCAAATTCC |
| nCoV 3,130 F pool 2 | GTGAAGAAGAAGAGTTTGAGCCA |
| nCoV 1,659 R pool 2 | GACCTTCGGAACCTTCTCCA |
| nCoV 1,510 F pool 1 | TTCTTCGTAAGGGTGGTCGC |
| nCoV 89 F pool 2 | ACCAACCAACTTTCGATCTCT |

**Data S1.** Metagenomics data obtained from respiratory specimens from Tiger 1 have been deposited in SRR11587605 under PRJNA627354.

**Data S2.** Whole genome consensus sequences obtained from SARS-CoV-2 detected in fecal samples from Tigers 2-4 and Lions 1-3 have been deposited in Gene Bank under accession numbers: MT704313, MT704315, MT704316, MT704312, MT704310, MT704311.

**Data S3.** Whole genome sequences obtained from SARS-CoV-2 tiger and lion isolates (TGR/NY-1/20, TGR/NY-3/20 and LN/NY-2/20) have been deposited in GenBank under accession numbers: MT704317, MT704314, MT747978.

**Data S4.** Whole genome sequences obtained from SARS-CoV-2 strains in Keepers 1 and 2 have been deposited in Gene Bank under accession numbers: MT703883, MT703884.
